# Earthworm and Soil Microbial Communities in Flower Strips

**DOI:** 10.1101/2022.12.15.520680

**Authors:** Zita Bednar, Anna Vaupel, Simon Blümel, Nadine Herwig, Bernd Hommel, Verena Haberlah-Korr, Lukas Beule

**Affiliations:** Julius Kühn Institute (JKI)–Federal Research Centre for Cultivated Plants, Institute for Ecological Chemistry, Plant Analysis and Stored Product Protection, Berlin, Germany; South Westphalia University of Applied Sciences, Department of Agriculture, Soest, Germany; Ruhr University Bochum, Faculty of Biology and Biotechnology, Bochum, Germany

**Keywords:** flower strips, earthworms, soil microbiome, soil archaea, soil bacteria, soil fungi, soil-N-cycling genes, arbuscular mycorrhizal fungi (AMF)

## Abstract

Flower strips are a common agricultural practice to increase aboveground biodiversity and beneficial ecosystem services. Although soil communities are a key component of terrestrial biodiversity and drive important ecosystem services, their abundance, diversity, and composition in flower strips remain largely unexplored. Here, we shed light on earthworms and soil microorganisms in flower strips and aim to provide a starting point for research on belowground communities in flower strips. In 2020, we established a field margin vegetation as well as two annual and two perennial flower strip mixtures at three study sites in Germany that were previously conventional croplands or fallow. Two years following this conversion, we determined earthworm communities and investigated the soil microbiome using real-time PCR (archaea, bacteria, fungi, and soil-N-cycling genes) and amplicon sequencing (bacteria and fungi). Different plant mixtures (i.e. field margin, annual, and perennial flower strips) harbored distinct earthworm and soil microbial communities. Earthworm density and biomass declined or remained unaffected in annual flower strips but increased in perennial flower strips as compared to field margins. Arbuscular mycorrhizal fungi showed greater diversity and community share in non-tilled (i.e. field margin and perennial flower strips) as compared to tilled plant mixtures (i.e. annual flower strips). We attribute changes in earthworms and microorganisms mainly to the effect of tillage and plant diversity. Overall, we suggest that perennial flower strips serve as refugia for soil biota in agricultural landscapes. Future studies should compare soil biota in perennial flower strips to those in adjacent fields and investigate whether beneficial belowground effects are restricted to the flower strips or spatially extend into adjacent fields (‘spillover’).

## Introduction

The global loss of biodiversity has far-reaching negative impacts on ecosystem functions [1] and consequently humanity [2]. Agricultural intensification significantly contributes to the loss of biodiversity in agroecosystems (e.g. [3]). Incorporation of flower strips along field edges is an established measure that is known to increase, maintain or restore biodiversity and its related ecosystem functions in agroecosystems. For example, flower strips provide habitat and food resources for pollinators and therefore promote their abundance and diversity (e.g. [4]). The magnitude of the effects of flower strips on pollination services and crop yield in adjacent croplands is variable and depends on the age of the flower strip and its plant diversity (i.e. perennial and old flower strips with high plant diversity promote pollination services most effectively) [5]. Furthermore, flower strips can increase the abundance of natural enemies of pests and promote pest control services (e.g. [6]). A recent data synthesis revealed that flower strips enhance pest control services in adjacent croplands by 16% on average [5].

Despite the large body of literature on aboveground biodiversity and ecosystem services, effects of flower strips on belowground communities remain largely unknown. Recently, it was postulated that flower strips impact soil biodiversity below them [7]. Considering the complex interactions between plants and soil biota as well as the impacts of agricultural management practices (e.g. tillage and crop rotation) on the abundance, community composition, and function of soil biota (e.g. [8]), we consider this assumption reasonable. Yet, experimental data on the effects of flower strips on soil biota are scarce in the scientific literature.

In this work, we shed light on soil biota under flower strips and aim to provide a starting point for research in this direction. For the first time, we investigated soil archaea, bacteria, fungi, and earthworms under a field margin vegetation versus four different types of flower strips (two annual and two perennial flower strip mixtures comprising 11 to 13 and 30 to 51 plant species, respectively) in three different soils. We hypothesized that i) flower strips increase the abundance and diversity of soil biota compared to field margin vegetation. We further expected that ii) perennial flower strips promote soil biota and their diversity more effectively than annual flower strips due to the absence of soil management (annual flower strips were re-establishment every spring) and larger plant diversity.

## Materials & Methods

### Study site and study design

Our study was conducted at three study sites (near Lippetal on a Gleyic Podzol, at the experimental research station of the South Westphalia University of Applied Sciences near Merklingsen on a Gleyic Luvisol, and near Ense on a Stagnic Cambisol; Fig 1; see S1 Table for site description and general soil properties) in the federal state of North Rhine-Westphalia, Germany. We refer to the study sites by their soil group (i.e. Podzol, Luvisol, and Cambisol soil).

**Fig 1.**
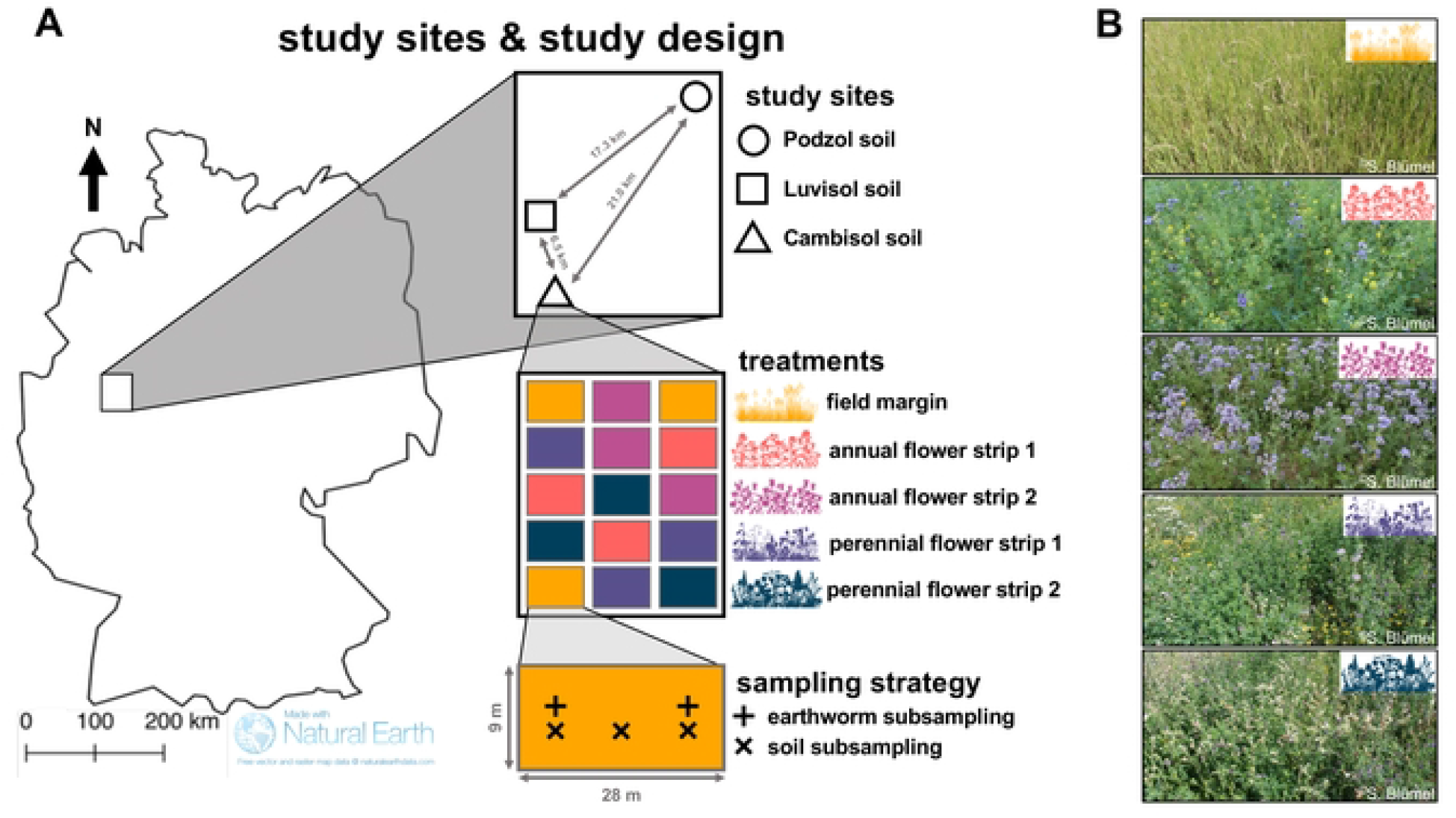
Study sites and study design. Study sites and study design (A) and photos of the flower strips taken in July 2022 at the study site on the Cambisol soil (B). Images are courtesy of the Integration and Application Network (ian.umces.edu/media-library).

In 2020, five different plant mixtures were established (the soil was tilled twice (grubber and rotary harrow) prior to sowing due to weed pressure) at a seeding rate of 10 kg ha^-1^ at each site. For each plant mixture, three replicate plots of 9 × 28 m were established at each site in a completely randomized design (3 study sites x 5 plant mixtures x 3 replicate plots = 45 replicate plots across sites) (Fig 1). A field margin vegetation was established in autumn 2020 by sowing a mixture of four grasses commonly found in field margins at our study region (referred to as ‘field margin’). Four different flower strip mixtures were established in spring 2020 using two annual flower strip mixtures (comprising 11 and 13 plant species, referred to as ‘annual flower strip 1’ and ‘annual flower strip 2’, respectively) and two perennial flower strip mixtures (comprising 30 and 51 plant species, referred to as ‘perennial flower strip 1’ and ‘perennial flower strip 2’, respectively) (Fig 1). The floral composition of the five different plant mixtures at sowing is given in S2 Table. Prior to the experiment, the sites were conventionally managed croplands (Podzol and Cambisol soil) or fallow (Luvisoil soil).

At each site, the annual flower strips were re-established (flower strips were mulched and the soil was tilled twice (grubber and rotary harrow) prior to resowing) in April 2021 and 2022. The field margin and perennial flower strips were topped at 15 cm height in March 2022 and not further managed, except in the Luvisol soil where all flower strips had to be re-established in spring 2021 due to high weed pressure. None of the replicate plots received fertilizer or plant protection products during the experiment.

### Soil sampling

Soil samples for the analysis of general soil properties (soil pH, organic C, total N, and bulk density) and soil microorganisms were collected from July 15 to 16 2022. Soil samples at 0 – 5 m soil depth were collected using a 250 cm^3^ stainless steel cylinder, whereas soil samples at 0 – 30 cm depth were collected using a stainless-steel auger (⌀ 3.5 cm). At each replicate plot, three soil subsamples were collected per depth and thoroughly homogenized in a sterile polyethylene bag to obtain one composite soil sample for each depth at each replicate plot. From the composite samples, an aliquot of approximately 50 g fresh soil was stored at −20°C in the field for molecular analysis of soil microbial communities. Upon arrival at the laboratory, frozen soil samples were stored at −20°C until freeze-drying.

### Determination of general soil properties

Soil bulk density was determined at 0 – 5 cm soil depth with 250 cm^3^ stainless steel cylinders using the soil core method [9]. Prior to determination of other soil properties, soil samples were air-dried and sieved to < 2 mm. Soil pH, soil organic C (SOC) and total N were measured at 0 – 5 and 0 – 30 cm soil depth. Double lactate-extractable P (P_DL_) and K (K_DL_), calcium chloride-extractable Mg (Mg_CaCl2_), and soil texture were measured at 0 – 30 cm soil depth. Soil pH was determined in demineralized H_2_O at a ratio of 1:2.5 (soil:water (w/v)). Prior to the determination of SOC, carbonates were removed from the samples using acid fumigation as per [10]. SOC and total N were determined using a CNS elemental analyzer (Vario EL Cube, Elementar, Germany). P_DL_ and K_DL_ were determined as per [11] and Mg_CaCl2_ as per [12]. Soil texture was determined as per [13].

### Earthworm extraction

Earthworm communities were sampled from October 16 to 18 2022 using Allyl isothiocyanate (AITC) expulsion as described previously [14]. Briefly, within each replicate plot, earthworms were expelled from two subplots in order to account for spatial heterogeneity. Squared aluminum frames (50 × 50 cm) were embedded approx. 5 cm into the soil and 5 liters of a 0.01% (v/v in tap water) AITC solution were poured into the frames. Emerging earthworms were collected from the soil surface for 30 minutes, washed with tap water, and stored in tap water. Within 12 hours post sampling, earthworms were weighted, species were determined based on morphology, and all collected individuals were released. Earthworm counts and biomass from the two subplots were added up. Earthworm species were classified into three ecological groups: epigeic, endogeic, and anecic earthworms.

### Soil DNA extraction

Frozen soil samples were freeze-dried for 72 hours and thoroughly homogenized using a vortexer as described previously [15]. DNA was extracted from 50 mg finely ground soil using a cetyltrimethylammonium bromide (CTAB)-based protocol as per [16]. Quantity and quality of the DNA extracts were assessed on 1.7% (w/v) agarose gels stained with SYBR Green I solution (Thermo Fisher Scientific GmbH, Dreieich, Germany).

### Quantification of soil microbial groups using real-time PCR

Prior to real-time PCR, DNA extracts were diluted 1:50 (v/v) in double distilled H_2_O (ddH_2_O) to overcome PCR inhibition [17]. Soil bacteria and fungi were quantified as described previously [18]. Soil archaea were quantified using the primer pair 340F / 100R[19] using the identical master mix composition as for fungi [18]. The thermocycling conditions of archaea were as follows: initial denaturation at 95°C for 120 sec followed by 40 cycles of 95°C for 20 sec, 60°C for 30 sec, and 68°C for 30 sec, and final elongation at 68°C for 5 min. Genes involved in soil nitrogen (N)-cycling (nitrification: ammonia-oxidizing archaea (AOA) and bacteria (AOB) *amoA* genes; denitrification: *nirK, nirS*, and *nosZ* clade I and II genes) were quantified to estimate the population size of N-cycling microorganisms as per [15]. All reactions were carried out in 4 µL reaction volumes in a Peqstar 96Q thermocycler (PEQLAB, Erlangen, Germany). Melting curves were generated as described previously [15].

### Amplicon sequencing of the soil microbiome

Soil bacteria and fungi were amplified using the primer pair 341F (5′-CCTACGGGNGGCWGCAG-3′) / 785R (5′-GACTACHVGGGTATCTAAKCC-3′ [20] and ITS1-F_KYO2 (5’-TAGAGGAAGTAAAAGTCGTAA-3’) [21] / ITS86R (5’-TTCAAAGATTCGATGATTCA-3’) [22], respectively. Prior to PCR, DNA extracts were diluted 1:50 (v/v) in ddH_2_O to overcome PCR inhibition [17]. Amplification was carried out in 25 µL reaction volume in an Eppendorf Mastercycler EP Gradient S thermocycler (Eppendorf, Hamburg, Germany). Bacteria and fungi were each amplified within one PCR run using the same mastermix for all samples. The reaction volume contained 18.75 µL mastermix and 6.25 µL template DNA or ddH_2_O for a negative control. The mastermix comprised ddH_2_O, buffer (10 mM Tris-HCl, 50 mM KCl, 2.0 mM MgCl_2_, pH 8.3 at 25°C), 100 µM of each deoxynucleoside triphosphate (New England Biolabs, Beverly, Massachusetts, USA), 0.5 µM of each primer, 1 mg mL^−^bovine serum albumin, and 0.03 u µL^−^ Hot Start *Taq* DNA Polymerase (New England Biolabs, Beverly, Massachusetts, USA). Each primer was a mixture of primer with (50%) and without (50%) Illumina TruSeq 5’-end adapters (5’-GACGTGTGCTCTTCCGATCT-3’ for the forward primer and 5’-ACACGACGCTCTTCCGATCT-3’ for the reverse primer). Bacteria and fungi were amplified using a touch-up PCR protocol [23] with initial denaturation at 95°C for 2 min, 3 touch-up cycles (95°C for 20 sec, 50°C for 30 sec, and 68°C for 60 sec), 22 or 25 cycles (95°C for 20 sec, 58°C for 30 sec, and 68°C for 60 sec) for bacteria and fungi, respectively, and final elongation at 68°C for 10 min. Amplification success was verified on 1.7% (w/v) agarose gel stained with SYBR Green I solution (Thermo Fisher Scientific GmbH, Dreieich, Germany) and libraries were shipped to LGC Genomics (Berlin, Germany). A second amplification with standard i7- and i5-sequencing adapters was performed at the facilities of LGC Genomics. Libraries were multiplexed and sequenced on an Illumina MiSeq (V3 chemistry, 2 × 300 bp) (Illumina, Inc., San Diego, CA, USA). Amplicon sequencing data have been deposited at NCBI’s Short Read Archive (BioProject https://dataview.ncbi.nlm.nih.gov/object/PRJNA905898?reviewer=nu62fi5608g31f2tc85ne4beun for bacteria and https://dataview.ncbi.nlm.nih.gov/object/PRJNA905904?reviewer=7h9takgoltgqkihf476vo3v92q for fungi).

### Bioinformatic processing of amplicon sequencing data

Paired-end sequencing data of bacteria and fungi were demultiplexed using Illumina’s bcl2fast version 2.20 (Illumina, San Diego, CA, USA). One-sided and conflicting barcodes as well as barcodes containing more than two mismatches were removed. Sequencing adapter and primer sequences were clipped and reads with < 100 bp were discarded. Afterwards, sequencing reads were processed in QIIME 2 version 2022.2 [24]. Quality scores were manually inspected using the ‘q2-demux’ plugin. Sequence reads were quality filtered (allowing two expected errors), merged, and cleaned from chimeric sequences and singletons using DADA2 [25]. Obtained amplicon sequencing variants (ASVs) of bacteria and fungi were taxonomically classified against the SILVA ribosomal RNA gene database version 138 [26] and UNITE database version 8.3 QIIME developer release [27], respectively. Classification was achieved utilizing a scikit-learn Naive Bayes machine-learning classifier (‘q2-fit-classifier-naive-bayes’ and ‘q2-classify-sklearn’ plugin) in the ‘balanced’ configuration ([7,7]; 0.7 for bacteria and [6,6]; 0.96 for fungi as suggested by [28]). Following classification, non-bacterial and non-fungal sequence reads were discarded from the bacterial and fungal data sets. Scaling with ranked subsampling (SRS) [29] using the ‘SRS’ R package version 0.2.3 [30] was used to normalize the bacterial and fungal ASV table to 19,219 and 18,318 sequence counts per sample, respectively. The normalized data sets contained 44,009 bacterial and 3,648 fungal ASVs.

### Statistical analysis

All data were manually inspected for homoscedasticity and normal distribution of the residuals and tested using Levene’s and Shapiro-Wilk test, respectively. Relative change of earthworm density and biomass as well as the abundance of archaea, bacteria, fungi, and N-cycling genes in response to the flower strips was calculated as follows:

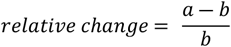

where *a* is the replicate plot of the flower strip mixtures or the field margin and *b* is the mean of the replicate plots of field margin per study site (i.e. soil type).

Alpha diversity indices (i.e. Shannon index (*H’*), Chao1 index, and Pielou’s evenness (*J’*)) of bacterial and fungal communities were determined using the ‘vegan’ R-package (version 2.5-7) [31]. Using the same R-package, pairwise Bray-Curtis dissimilarities were calculated and visualized using non-metric multidimensional scaling (NMDS). Additionally, differences in community composition were determined using permutational multivariate analysis of variance (PERMANOVA) on Bray-Curtis dissimilarities using 999 permutations. PERMANOVA was run to test the effects of study site (i.e. soil type) and plant mixture (i.e. field margin and different flower strips) [adonis2(dissimilarity matrix ∼ soil type + plant mixture + soil type:plant mixture, nperm = 999)] on the community composition. Additionally, we tested the effect of plant mixture within each soil type. Complementary to each PERMANOVA model, we assessed the dispersion of samples in each group using multivariate homogeneity of group dispersions.

Differences in earthworm density and biomass, absolute abundance of archaea, bacteria, fungi, and N-cycling genes, alpha diversity indices of bacteria and fungi, and soil properties were determined using one-way analysis of variance (ANOVA). Differences in relative abundance of taxa were determined from log(x+1)-transformed data. Correlations among different parameters were performed using Spearman rank correlations. All statistical analyses were performed in R (version 4.1.2) [32]. For all statistical test, statistical significance was considered at p < 0.05.

## Results

### General soil properties

Within each soil type, soil properties remained unaffected by the recent introduction of flower strips. Flower strips did not affect soil pH, bulk density, SOC, total N, P_DL_, K_DL_, and Mg_CaCl2_.

### Earthworm communities

Earthworm density and biomass were strongly correlated (*r* = 0.95; p < 0.0001) and increased from the Podzol to the Luvisol to the Cambisol soil (Fig 2 A, S1 Fig). Seven different earthworm species were found across the three study sites: *Allolobophora chlorotica, Aporrectodea caliginosa, Aporrectodea longa, Aporrectodea rosea, Aporrectodea trapezoides* (also referred to as a subspecies of *Aporrectodea caliginosa*), *Lumbricus rubellus*, and *Lumbricus terrestris*. The classification of the species into ecological groups (i.e. anecic, endogeic, and epigeic) revealed that earthworm community composition was site-specific. In the Podzol soil, anecic earthworms were absent and epigeic earthworms accounted for a large share of the community. In contrast, epigeic earthworms were not present in the Luvisol soil. The Cambisol soil harbored all three ecological groups (Fig 2 B).

**Fig 2.**
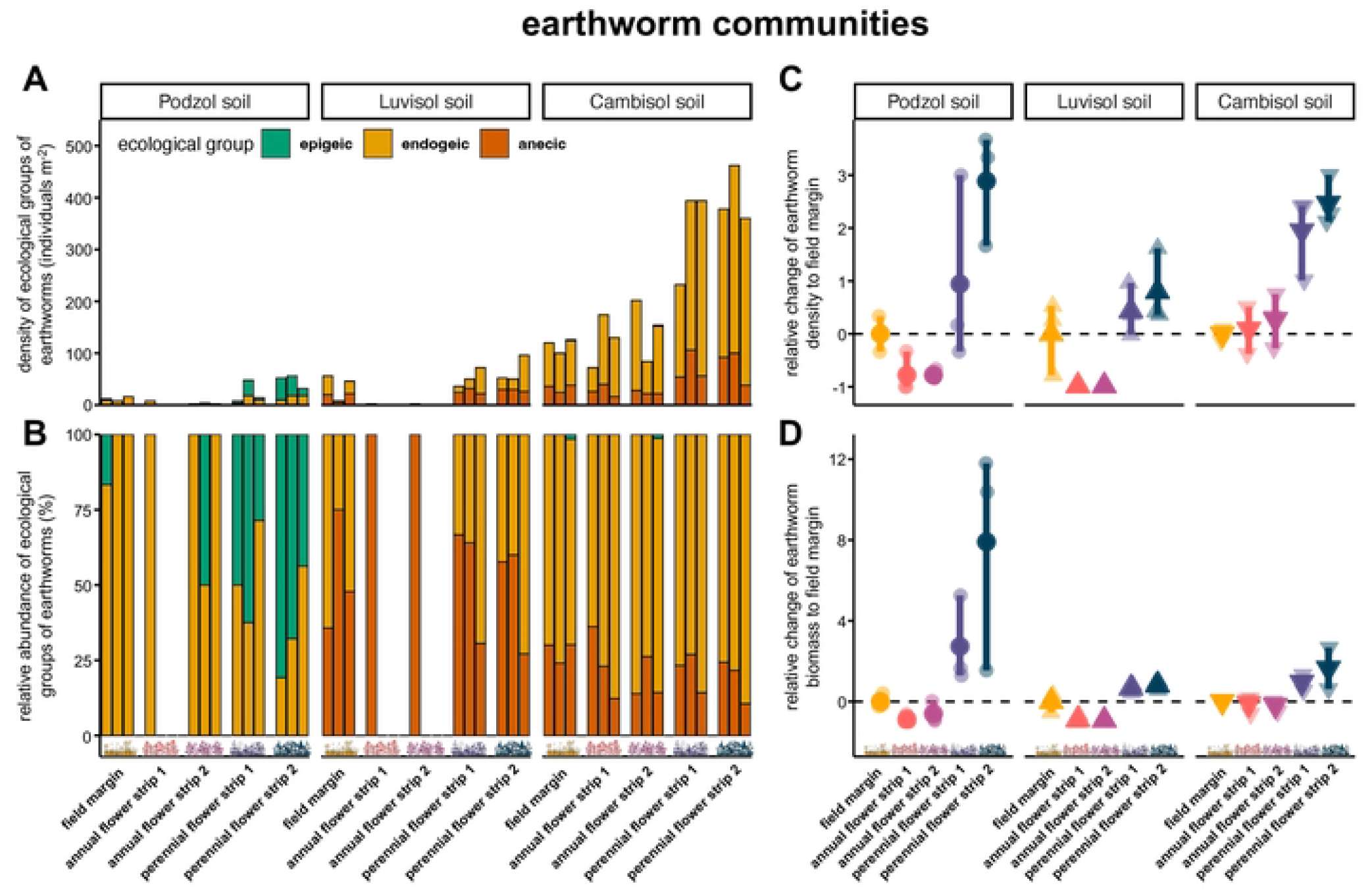
Earthworm communities. Population densities of ecological groups of earthworms (**A**) and their relative abundance within the earthworm communities (**B**). Bars represent individual replicate plots (*n =* 3). Relative change of earthworm density (**C**) and biomass (**D**) in response to flower strips. Non-transparent dots and triangles represent means and vertical bars represent standard deviation. Transparent dots and triangles represent individual data points (i.e. replicate plots). Images are courtesy of the Integration and Application Network (ian.umces.edu/media-library).

Perennial flower strips strongly promoted earthworm population density and biomass across soils (Fig 2 A, C, D). In contrast, annual flower strips showed consistently lower density and biomass than the field margin in the Podzol and Luvisol soil (Fig 2 A, C, D). In these two soils, earthworms were almost absent under the annual flower strips (Fig 2 A). In the Podzol soil, the perennial flower strip 2 increased earthworm density and biomass by a factor of 3.7 to 17.5 compared to the field margin and the annual flower strips (p 2264 0.031), which was mainly driven by the increased occurrence of epigeic earthworms in the perennial flower strip 2. Earthworm density in the flower strips in the Luvisol soil did not differ statistically significant from the field margin. However, earthworm densities were 79 to 99 times larger in perennial than in annual flower strips (p ≤ 0.036). In the same soil, earthworm biomass was 15.4 to 23.3 times larger in perennial flower strips and 9.2 to 12.8 times larger in the field margin (p ≤ 0.025) as compared to annual flower strips. The Cambisol soil was the only soil in which annual flower strips showed earthworm densities and biomass similar to those in the field margin. In this soil, perennial flower strips increased earthworm density by 171 to 247% as compared to the annual flower strips and field margin (p ≤ 0.018); earthworm species richness remained unchanged.

### Soil microbiome

Population sizes of archaea, bacteria, fungi, and functional groups involved in soil N-cycling were not affected by flower strips (S2 Fig, S3). Across soils, soil bacterial communities were dominated by the phyla of *Actinobacteriota* (29.4 ± 6.1%), *Proteobacteria* (16.4 ± 1.8%), and *Acidobacteriota* (12.5 ± 1.8%). The dominating bacterial classes were *Actinobacteria* (20.1 ± 6.6%), *Alphaproteobacteria* (11.3 ± 1.1%), and *Planctomycetes* (7.9 ± 2.3%) (Fig 3 A). The fungal community was dominated by *Ascomycota* (65.4 ± 14.0%), *Mortierellomycota* (12.3 ± 9.5%), and *Basidiomycota* (12.0 ± 8.2%) on phylum level and *Sordariomycetes* (41.5 ± 16.0%), *Dothideomycetes* (17.2 ± 11. 6%), and *Mortierellomycetes* (12.2 ± 8.1%) on class level (Fig 3 E). Alpha diversity indices (Shannon index (*H’*), Chao1 index, and Pielou’s evenness (*J’*)) were not affected by flower strips (Fig 3 B, C, D, F, G, H) except fungal Shannon diversity in the Luvisol soil which was higher in the perennial flower strips and the field margin compared to the annual flower strip 2 (p = 0.036) (Fig 3 F).

**Fig 3.**
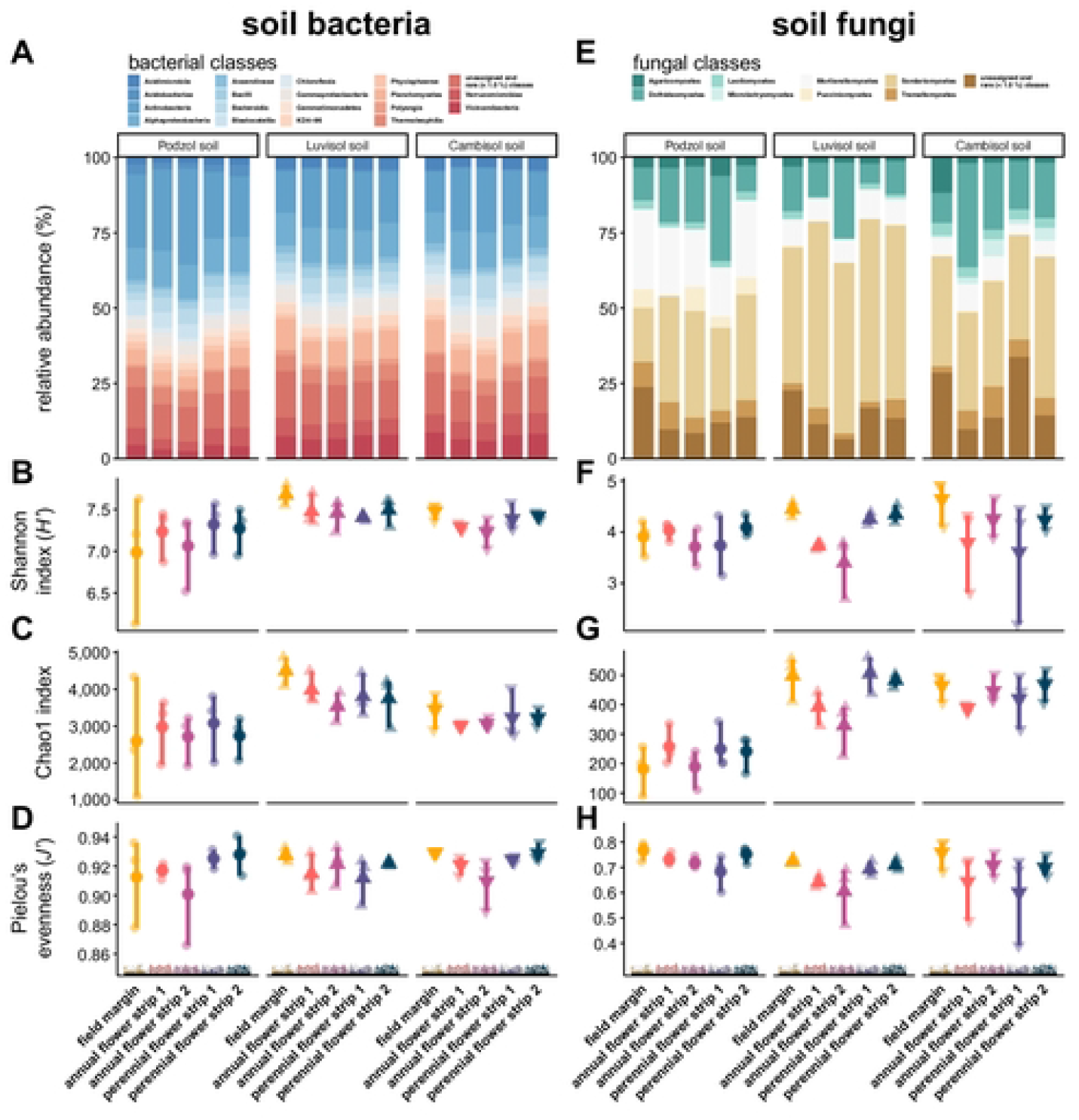
Community composition and alpha diversity of soil bacteria and fungi. Mean relative abundance of bacterial (**A**) and fungal classes (**B**) per plant mixture and soil type. Alpha diversity indices of bacterial (**B, C, D**) and fungal communities (**F, G, H**). Non-transparent dots and triangles represent means and vertical bars represent standard error (*n =* 3). Transparent dots and triangles represent individual data points (i.e. replicate plots). Images are courtesy of the Integration and Application Network (ian.umces.edu/media-library).

Soil type (i.e. Podzol, Luvisol, and Cambisol soil) and plant mixture (i.e. field margin and different flower strips) affected community composition of both bacteria and fungi (Table 1). For both communities, the effect of soil type on community composition was stronger than the effect of plant mixture (Table 1). Plant mixture effects per site were visualized using NMDS (Fig 4). In the Luvisol and Cambisol soil the field margin, the annual flower strips, and the perennial flower strips each formed a distinct cluster in the NMDS for both bacteria and fungi (Fig 4 B, C, E, F). In the Podzol soil, two clusters emerged comprising the non-tilled plant mixtures (i.e. the field margin and the perennial flower strips) and the tilled plant mixtures (i.e. the annual flower strips) (Fig 4 A). In the same soil, soil fungal communities per plant mixture clustered almost separately (Fig 4 B).

**Table 1.**
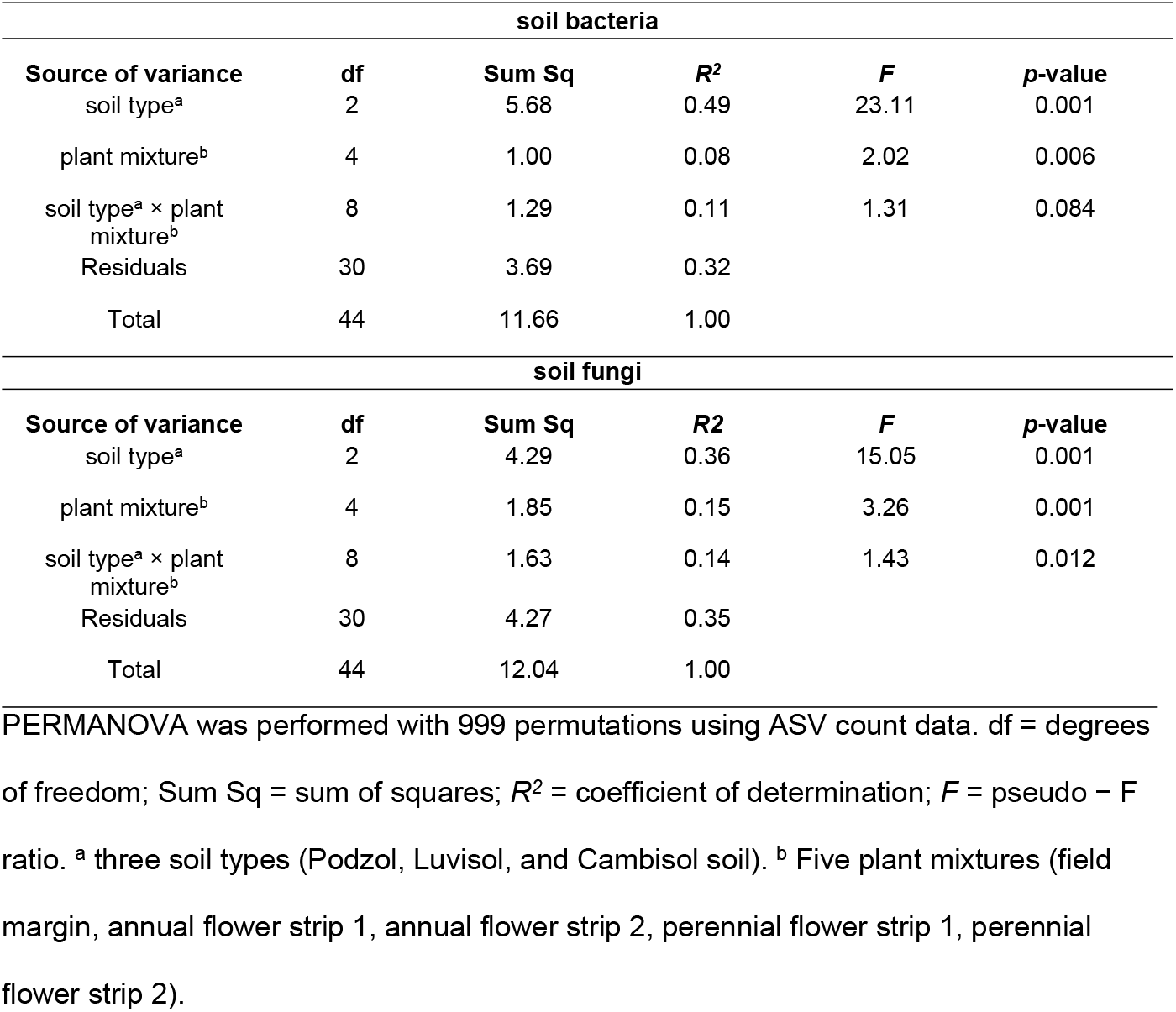
Permutational multivariate analysis of variance (PERMANOVA) results for soil bacterial and fungal community composition.

**Fig 4.**
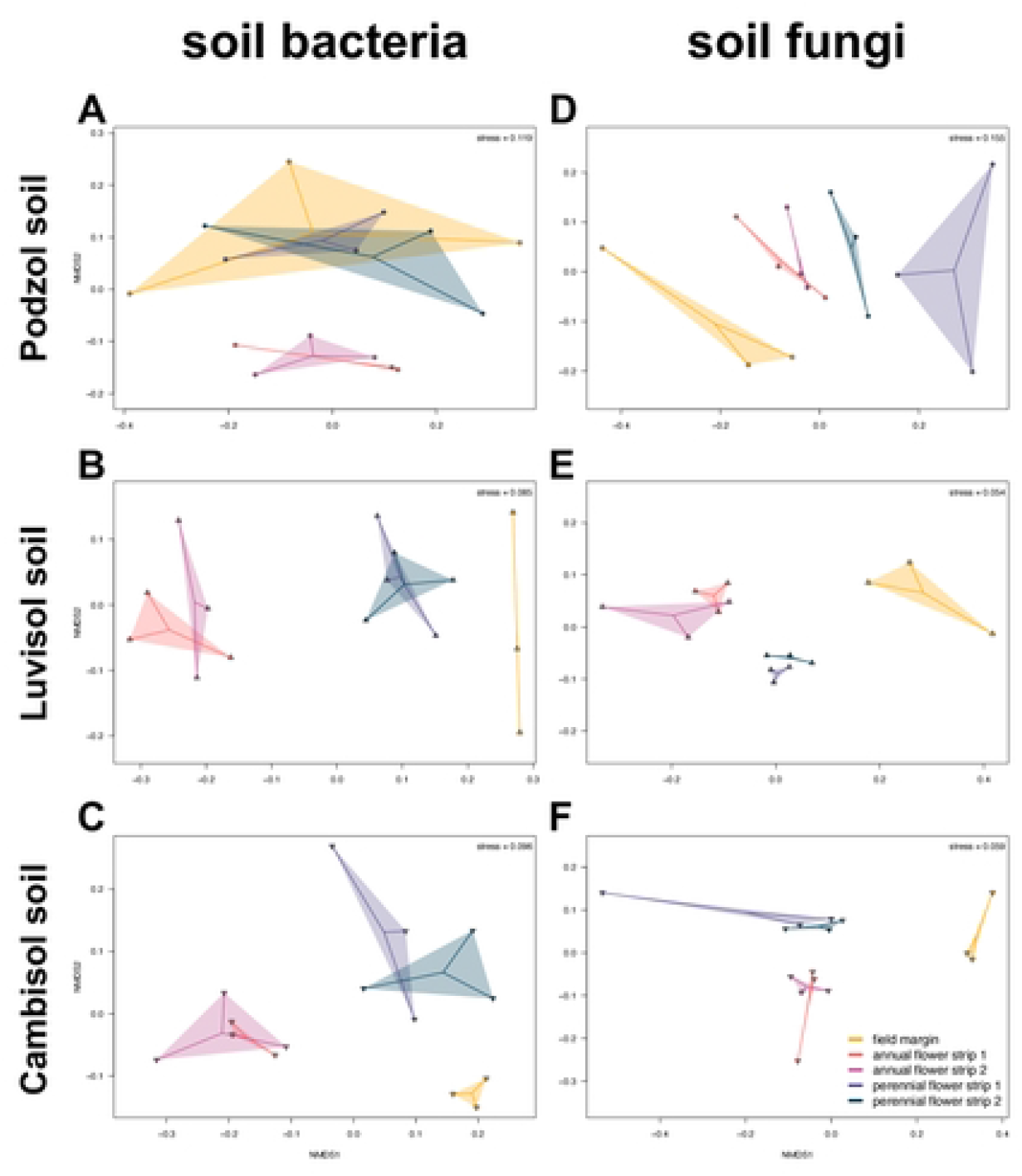
Non-metric multidimensional scaling (NMDS) of Bray-Curtis dissimilarities of soil bacterial and fungal communities. NMDS plots of bacterial (**A, B, C**) and fungal communities (**D, E, F**) within each soil type. Dots and triangles represent individual data points (i.e. replicate plots) (*n =* 3) which are connected with the centroid of their respective plant mixture.

The community share of several bacterial phyla was affected by the plant mixtures (Fig 5; see S3 Table for p-values) and reflected the clustering in the NMDS. For example, relative abundance of *Desulfobacterota* in the Cambisol soil were greater in the field margin than in the flower strips (p ≤ 0.0001). Similarly, in the Luvisol soil, relative abundance of *Methylomirabilota, NB1-j, Planctomycetota* was greater in the field margin as compared to the annual flower strips (p ≤ 0.015). In the same soil, *Latescibacterota* showed greater relative abundance in the field margin than in the flower strips (p ≤ 0.0011). Additionally, relative abundance of *Latescibacterota* was greater in the perennial than in the annual flower strips (p ≤ 0.046). In the Cambisol soil, the field margin increased the relative abundance of *Methylomirabilota* and *Latescibacterota* as compared to the annual flower strips (p ≤ 0.015). In the same soil, *Planctomycetota* showed greater relative abundance in the field margin and perennial flower strips than in the annual flower strips (p ≤ 0.022). Compared to the field margin, annual flower strips promoted the relative abundance of *Actinobacteria, Bdellovibrionota*, and *Proteobacteria* in the Cambisol soil (p ≤ 0.037). Likewise, in the Luvisol soil, relative abundances of *Abditibacteriota* and *Gemmatimonadota* were greater in the annual flower strips than in the field margin (p ≤ 0.043). In the same soil, relative abundance of *Bdellovibrionota* was greater in the annual flower strips as compared to the field margin and perennial flower strips (p ≤ 0.0048). In all soil types, relative abundance of *Bacteroidota* were greater in the annual flower strips than in the field margin (p ≤ 0.021).

**Fig 5.**
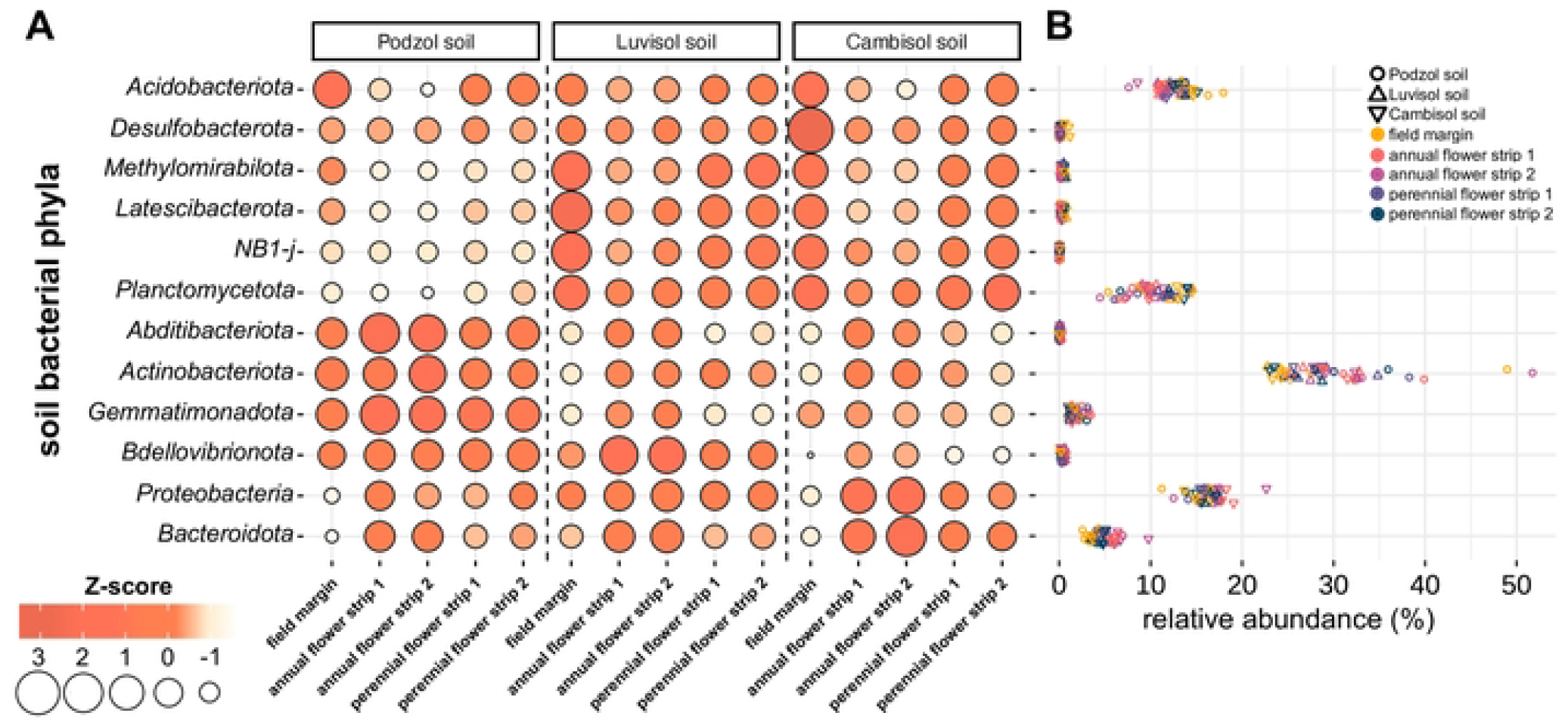
Selected soil bacterial phyla in flower strips. Z-score normalized relative abundance (**A**) and relative abundance (**B**) of bacterial phyla in three different soil types. Colored dots and triangles represent individual data points (i.e. replicate plots) (**B**).

Within the fungal community, the abundance and diversity of affiliates of the monophyletic phylum *Glomeromycota* (containing all arbuscular mycorrhizal fungi (AMF)), were altered by the plant mixtures (Fig 6). In all soil types, relative abundance of AMF was greater in the field margin as compared to the annual flower strips (p ≤ 0.007). Furthermore, in the Cambisol soil, relative abundance of AMF in the perennial flower strips was lower than in the field margin (p ≤ 0.0003), whereas in the Podzol soil, this was true only for the perennial flower strip 1 (p ≤ 0.032). In the Luvisol soil, the relative abundance of AMF in the field margin was greater than in the annual flower strips (p ≤ 0.0005) as well as in the perennial flower strip 1 (p ≤ 0.016). In the perennial flower strip 2, the abundance of AMF was greater compared to the annual flower strip 1 (p ≤ 0.042) but lower than in the field margin (p ≤ 0.049).

**Fig 6.**
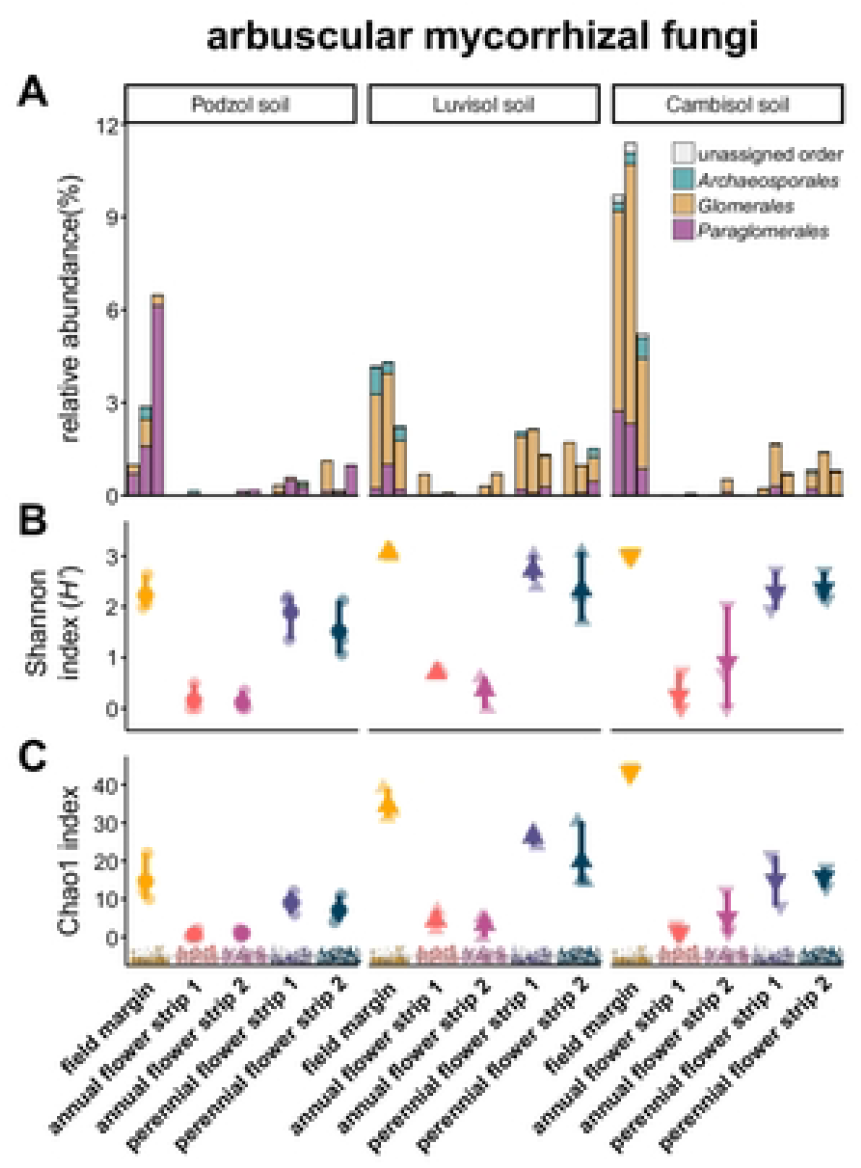
Arbuscular mycorrhizal fungi (AMF – Glomeromycota) in flower strips. Relative abundance of AMF orders in three different soil types (**A**). Bars represent individual replicate plots (*n =* 3). Shannon (*H’*) (**B**) and Chao1 index (**C**) of AMF. Non-transparent dots and triangles represent means and vertical bars represent standard error (*n =* 3). Transparent dots and triangles represent individual data points (i.e. replicate plots). Images are courtesy of the Integration and Application Network (ian.umces.edu/media-library).

Across sites and plant mixtures, 249 ASVs were assigned to AMF, covering three orders, namely *Archaeosporales, Glomerales*, and *Paraglomerales* (Fig 6 A). Relative abundance of *Archaeosporales* in the Luvisol and the Cambisol soil was greater in the field margin as compared to the annual and perennial flower strips (p ≤ 0.016). Furthermore, in the Luvisol soil, relative abundance of *Glomerales* was greater in the field margin and perennial flower strip 1 than in the annual flower strips (p ≤ 0.043). Relative abundance of *Glomerales* in the Cambisol soil was greater in the field margin compared to the annual and perennial flower strips (p ≤ 0.0008). In the Podzol soil, plant mixtures only affected the community share of *Paraglomerales* which was greater in the field margin compared to the annual flower strips (p ≤ 0.036). In the Cambisol soil, relative abundance of Paraglomerales was lower in the annual and perennial flower strips as compared to the field margin (p ≤ 0.0007). In contrast, community share of *Paraglomerales* did not differ among plant mixtures in the Luvisol soil.

Alpha diversity (Shannon index (*H’*) and Chao1 index) of AMF differed significantly among the plant mixtures (Fig 6 B, C). In each soil type, alpha diversity of AMF was greater in the field margin as compared to the annual flower strips (p ≤ 0.0053 and p ≤ 0.0066 for Shannon index and Chao1, respectively). Furthermore, alpha diversity of AMF did not differ between field margin and the perennial flower strips in the Podzol and Luvisol soil. In the Cambisol soil, however, Chao1 index was greater in the field margin than in the perennial flower strips (p ≤ 0.0001). According to Shannon index, alpha diversity of AMF was greater in the perennial flower strips as compared to the annual flower strip 1 in all soil types (p ≤ 0.012). In contrast, Chao1 index of AMF did not differ between the annual and perennial flower strips, except in the Cambisol soil where Chao1 index was greater in the perennial flower strips compared to annual flower strip 1 (p ≤ 0.021).

## Discussion

The integration of flower strips in agroecosystems is a common practice in many regions of the temperate zone to increase aboveground biodiversity and enhance beneficial ecosystem services. Although soil communities are a key component of terrestrial biodiversity and their diversity and composition determine ecosystem multifunctionality [33], soil biota in flower strips remain largely unexplored.

### Earthworm communities

In their role as ecosystem engineers, earthworms contribute to several beneficial soil functions (e.g. water infiltration (e.g. [34]), suppression of phytopathogens (e.g. [35]), and cycling of nutrients (e.g. [36]) and enhance soil fertility (e.g. [37]). Overall, earthworms are suitable biological indicators for sustainable soil management in agriculture [38]. More than two decades ago, [39] conducted one of the first studies on soil biota in flower strips. The authors showed that conversion of a maize field into a wild flower strip increased the abundance of earthworms already after one year and reached a plateau after two years [39]. Although their results are reasonable, the authors did not investigate earthworm populations in the maize field throughout the entire course of the experiment which would have been needed to exclude the effects of potential seasonal fluctuations that may have influenced the results.

In the present study, croplands or fallow were converted into either a field margin, annual flower strips or perennial flower strips. In the Podzol and Luvisol soil, annual flower strips showed the lowest earthworm density and biomass (Fig 2 A, S1 Fig), which we attribute to their annual re-establishment that included tillage (grubber and rotary harrow). Tillage is well-known to affect density, biomass, and community composition of earthworms [40,41]. While density of anecic species generally decreases under tillage due to physical damage and the removal of plant litter from the soil surface (e.g. [41]), responses of endogeic species to tillage are rather inconsistent. While some studies showed that the density of endogeic species is either unaffected (e.g. [42,43]) or increased through tillage (e.g. [41,44]) due to the incorporation of plant residues that serve as a food resource, other studies found a negative impact of tillage on endogeic earthworm density (e.g. [45,46]). In view of these inconsistent results, [47] recently conducted a global meta-analysis on the effects of tillage on earthworm abundance and biomass. Their results revealed that the population densities of all three ecological groups benefit from reduced tillage and that epigeic and anecic species benefit more than endogeic [47]. Their results agree with our findings of a decline in all three ecological groups of earthworms (epigeic, endogeic, and anecic) under the tilled annual flower strips as compared to the non-tilled field margin and perennial flower strips (Fig 2 A).

Although differences in tillage regimes can explain the low earthworm densities in the annual flower strips, they do not explain the increased population densities in the non-tilled perennial flower strips as compared to the non-tilled field margin (Fig 2 A, C, D). The impacts of plant diversity and biomass on earthworm communities have frequently been studied in grasslands. While some studies revealed a positive impact of plant diversity and biomass on earthworm density and biomass [48–50], other studies were not able to confirm this [51,52]. These discrepancies among studies may be related to, *inter alia*, interactions with other soil biota [53] and plant community composition [53–56]. In our study, higher plant diversity in the perennial flower strips as compared to the field margin promoted earthworm density and biomass in all three soil types (Fig 2 A, C, D, S1 Fig). Although plant biomass was not determined in our study, previous studies showed that plant biomass production (and consequently plant litter production) generally increases with plant diversity (e.g. [57]). Thus, we suggest that compared to the field margin, earthworm communities in the perennial flower strips benefited from higher quantities of above- and belowground plant litter (i.e. food resources). We further suggest that perennial flower strips not just increase the quantity of food input but also alter its quality which may be even more important for soil decomposer communities (e.g. [53,58]).

### Soil microbiome

Soil type strongly affected community composition of both bacteria and fungi (Table 1) which was expected considering the strong influence of soil properties on soil microbial community composition [59–62]. For example, soil pH has been studied extensively as a predictor for the community composition of bacteria and fungi across various spatial scales. Several studies concluded that bacterial communities are generally more affected by soil pH than fungal (e.g. [59,60]) which is likely due to a wider range of pH optima for fungal growth [59].

In addition to soil type, plant mixture (i.e. field margin and different flower strips) was also identified as a determining factor of bacterial and fungal community composition (Table 1, Fig 4). Dissimilarities in community composition of bacteria and fungi between the annual flower strips and the other plant mixtures in each soil type (Fig 4) may be related to tillage during the re-establishment of the annual flower strips. There is compiling evidence of not only changes in microbial population size [61] but also in community composition of bacteria and fungi in response to tillage intensity (e.g. [62–65]). In light of the strong impact of tillage on soil structure [66] and the subsequent consequences for soil as a biological habitat [67], it is conclusive that tillage can affect the composition of the soil microbiome.

Besides differences in soil management, differences in plant species composition as well as diversity of the plant mixtures (field margin < annual flower strips < perennial flower strips) likely contributed to the observed changes in community composition. Considering the plant diversity, this assumption is supported by the differences in community composition between the non-tilled field margin and the non-tilled perennial flower strips. There are numerous interactions between plants and soil microorganisms that shape the soil microbiome. For example, plant root exudates shape the soil microbiome (especially in the rhizosphere) by recruiting plant-beneficial microorganisms [68]. The quantity and quality of root exudates depend on abiotic and biotic stressors but also plant species and age [69].Thus, it is reasonable to assume that microbial community composition was driven by the variation in the root exudation due to differences in plant species composition of the different plant mixtures. Indeed, a recent microcosm experiment proposed root exudates as an important link between plant diversity and soil microorganisms [70]. Furthermore, differences in plant species composition are expected to result in differences in the quantity and quality of above- (leaves, stalks) and belowground (roots) plant litter among plant mixtures which have been identified as a driver of microbial communities (e.g. [71]) and could thus have contributed to the observed community shifts.

The soil bacterial community composition was strongly affected by the plant mixture at phylum level (Fig 5). In agreement with previous studies (e.g. [72]), we suggest that such alterations in community composition are expected to result in altered microbiome functionality. There are several tools to predict functional potential profiles from the taxonomical profiles of microbiome data sets [73]; however, we decided to not use these tools because microbiome data generated from short-read amplicons may not be suitable to accurately predict microbiome functions [74]. Instead, we suggest that future studies should measure actual microbial processes in flower strips and link these with microbiome data in order to test whether flower strips alter the functionality of the soil microbiome. Although not affected by the plant mixtures, our quantification of genes involved in soil-N cycling (S3 Fig) is an initial step towards understanding microbial functions in flower strips.

In contrast to the differences in beta diversity (i.e. compositional dissimilarities among plant mixtures) discussed above, overall alpha diversity of bacteria and fungi remained mostly unaffected by the plant mixtures (Fig 3). These results agree with the findings of [75] who found that plant diversity in grasslands is a predictor of beta but not alpha diversity. Alpha diversity of AMF, however, was affected by the plant mixtures (Fig 6 B, C). In addition to the diversity of AMF, plant mixtures also affected the relative abundance of AMF (Fig 6 A). AMF form symbiotic associations with the majority of terrestrial plants and, *inter alia*, enhance nutrient acquisition by associated plants (e.g. [76]). Therefore, AMF recently received increasing attention for use as biofertilizers in sustainable agriculture [77]. The greater community share and diversity of AMF in the non-tilled (field margin and perennial flower strips) than in the tilled (annual flower strips) plant mixtures (Fig 6) agrees with previous studies that showed that reduced tillage favors AMF (e.g. [78,79]). Recently, [80] compared AMF communities in field margins to those in arable land and found that field margins alter AMF community composition and increase AMF diversity as compared to arable land. Few years earlier, [81] proposed that AMF colonization could take place via different nearby landscape elements such as field margins. Although neighboring croplands were not investigated in this study, we hypothesize that perennial flower strips serve as a reservoir for AMF and enhance AMF colonization of neighboring crops.

## Conclusion

Field margins, annual, and perennial flower strips harbor distinct earthworm and soil microbial communities. Compared to field margins, earthworm density and biomass declined or remained unaffected in annual flower strips but increased in perennial flower strips. Soil type was the strongest predictor of bacterial and fungal community composition. However, plant mixture (i.e. field margin, annual, and perennial flower strips) affected microbiome assembly within each soil type. Although overall alpha diversity of bacteria and fungi remained mostly unaffected by the plant mixtures, AMF showed greater diversity and community share in non-tilled (i.e. field margin and perennial flower strips) as compared to tilled plant mixtures (i.e. annual flower strips). We attribute the observed changes in soil biota mainly to differences in tillage and plant diversity. Overall, our data suggests that perennial flower strips serve as refugia for soil biota in agricultural landscapes. Thus, future studies should compare the population size, diversity, and functionality of soil biota in flower strips to those in adjacent agricultural fields in order to assess the belowground benefits of flower strips. Furthermore, we suggest to investigate whether beneficial effects on belowground biota are restricted to the perennial flower strips or spatially extend into adjacent agricultural fields (‘spillover’) as they do for certain aboveground biota. We hope that our work provides a starting point for research on the biodiversity and function of belowground communities in flower strips.

## Acknowledgments

The authors would like to thank Josef Beule for participating in soil sampling.

## Supporting Information Captions

**S1 Fig. Earthworm biomass**. Biomass (g m^-2^) of ecological groups of earthworms. Bars represent individual replicate plots (*n =* 3).

**S2 Fig. Relative change of (A) soil archaea, (B) bacteria, and (C) fungi in response to flower strips**. Non-transparent dots and triangles represent means and vertical bars represent standard deviation (*n =* 3). Transparent dots and triangles represent individual data points (i.e. replicate plots). Archaea, bacteria, and fungi were quantified by using real-time PCR (see *Quantification of soil microbial groups using real-time PCR* for details). See *Statistical analysis* for details regarding the calculation of the relative change. Images are courtesy of the Integration and Application Network (ian.umces.edu/media-library).

**S3 Fig. Relative change of ammonia-oxidizing archaea (AOA) *amoA* (A), *nirS* (B), *nosZ* clade I (C), and *nosZ* clade II genes (D) in response to flower strips**. Non-transparent dots and triangles represent means and vertical bars represent standard deviation (*n =* 3). Transparent dots and triangles represent individual data points (i.e. replicate plots). AOA *amoA, nirS*, and *nosZ* clade I and II genes were quantified by using real-time PCR (see *Quantification of soil microbial groups using real-time PCR* for details). See *Statistical analysis* for details regarding the calculation of the relative change. Images are courtesy of the Integration and Application Network (ian.umces.edu/media-library).

**S1 Table. Study site description and general soil properties**.

**S2 Table. Composition of the plant mixtures at sowing**.

**S3 Table. Mean ± standard deviation of the relative abundance of soil bacterial phyla (*n =* 3)**. Different uppercase letters of the same font indicate statistically significant differences (p < 0.05).

